# Keratin-Based Epidermal Green Autofluorescence is a Common Biomarker of Organ Injury

**DOI:** 10.1101/564112

**Authors:** Mingchao Zhang, Yujia Li, Jiucun Wang, Huiru Tang, Zhong Yang, Danhong Wu, Yue Tao, Hao He, Sijia Wang, Xingdong Chen, Shankai Yin, Haibo Shi, Xunbin Wei, Tianqing Chu, Wanzhi Tang, Dhruba Tara Maharjan, Zhaoxia Yang, Yu Wang, Li Jin, Weihai Ying

## Abstract

It is critical to discover biomarkers for non-invasive evaluation of the levels of inflammation and oxidative stress in human body - two key pathological factors in numerous diseases. Our study has indicated keratin 1-based epidermal autofluorescence (AF) as a biomarker of this type: Inducers of both inflammation and oxidative stress dose-dependently increased epidermal green AF with polyhedral structure in mice, with the AF intensity being highly associated with the dosages of the inducers. Lung cancer also induced increased epidermal green AF of mice, which was mediated by inflammation. Significant and asymmetrical increases in green AF intensity with polyhedral structure were found in the Dorsal Index Fingers’ skin of acute ischemic stroke (AIS) patients. While the AF intensity of the subjects with high risk for developing AIS, ischemic stroke patients in recovery phase and lung cancer patients was significantly higher than that of healthy controls, both AF intensity and AF asymmetry of these four groups were markedly lower than those of the AIS patients, which have shown promise for AIS diagnosis. Several lines of evidence have indicated K1 as an origin of the AF, e.g., K1 siRNA administration attenuated the oxidative stress-induced AF increase of mice. Collectively, our study has indicated K1-based epidermal AF as a biomarker for non-invasive evaluation of the levels of inflammation and oxidative stress in the body. These findings have established a basis for novel keratin’s AF-based biomedical imaging technology for non-invasive, efficient and economic diagnosis and screening of such inflammation- and oxidative stress-associated diseases as AIS.

## Introduction

Inflammation and oxidative stress are two key common pathological factors of multiple major diseases such as acute ischemic stroke (AIS)^1-4^. Chronic systemic inflammation and oxidative stress are closely related, which can promote development of a number of diseases such as cancer^5-7^. However, there has been no practical biomarker or method that can be used for determining non-invasively the levels of inflammation and oxidative stress in human body. Therefore, it is of critical significance to search for both biomarkers and methods of this type.

Human autofluorescence (AF) from the advanced glycation end-product (AGE)-modified collagen and elastin of dermis has shown promise for non-invasive diagnosis of diabetes and diabetes-related pathology^8^. Epidermal AF may also become a novel endogenous reporter of certain pathological insults, if the pathological insults can induce changes of epidermal AF. Keratins, melanin, NADH and FAD are four known epidermal fluorophores^9,10^. There are increased inflammation and reactive oxygen species (ROS) in impaired organs and other parts of the body such as the blood circulation in a number of diseases^11-16^. Therefore, we hypothesized that inflammation and ROS in the body may lead to changes of epidermal AF by affecting certain epidermal fluorophores, which may become endogenous reporters of the levels of inflammation and ROS in the body.

Keratins play multiple biological roles in epithelium, including intermediate filament formation^17^ and inflammatory responses^18^. Keratins have also been used as a tumor biomarker^19^. Keratin 1 (K1) / keratin 10 (K10) heterodimer is a hallmark for keratinocyte differentiation^20^, which is localized selectively in suprabasal differentiated keratinocytes^21^. However, there has been little information regarding the biomedical value of keratin’s AF.

In our current study, we tested the hypothesis regarding epidermal AF. Our study has indicated that K1-based epidermal AF is a biomarker of this type, which forms a basis for novel, keratin-based biomedical imaging technology for non-invasive evaluation of the levels of inflammation and oxidative stress in human body.

## Materials and Methods

### Materials

All chemicals were purchased from Sigma (St. Louis, MO, USA) except where noted. Male C57BL/6 mice, ICR mice, and BALB/cASlac-nu nude mice of SPF grade were purchased from SLRC Laboratory (Shanghai, China).

### Studies on Human Subjects

The studies on the group of AIS patients (AIS Group), the group of ischemic stroke patients in recovery phase (Recovery Group), the group of the persons with high risk for developing AIS (High-Risk Group), the group of lung cancer patients (Lung Cancer Group), and the group of healthy controls (Healthy Group) were conducted according to protocols approved by the Ethics Committee of Shanghai Fifth People’s Hospital Affiliated to Fudan University and the Ethics Committee of Shanghai Chest Hospital Affiliated to Shanghai Jiao Tong University. The AIS patients were the human subjects who were diagnosed as AIS patients and hospitalized in the Department of Neurology, Shanghai Fifth People’s Hospital Affiliated to Fudan University. The lung cancer patients were the human subjects who were diagnosed as lung cancer patients and hospitalized in the Department of Pulmonary Medicine, Shanghai Chest Hospital Affiliated to Shanghai Jiao Tong University. The human subjects of the High-Risk Group were hospitalized or evaluated in the Outpatient Clinic of the Department of Neurology, Shanghai Fifth People’s Hospital Affiliated to Fudan University. The subjects of the Recovery Group were the ischemic stroke patients who were evaluated in the Outpatient Clinic of the Department of Neurology, Shanghai Fifth People’s Hospital Affiliated to Fudan University one to three months after they had AIS. The average age of the Healthy Group, the High-Risk Group, the AIS Group, the Recovery Group, and the Lung Cancer Group was 63.65±0.92, 67.93±0.67, 64.11±1.02, 65.42 ±1.14 and 63.09±0.83 years of old, respectively.

### Development of a mouse model of lung cancer

Animal studies were conducted according to an animal protocol approved by the Animal Protocol Committee, School of Biomedical Engineering, Shanghai Jiao Tong University. Cell suspension of LLC (Lewis Lung Carcinoma) cells (1 × 10^6^ cells) in a total volume of 5 ml mixed with Matrigel (PBS : Matrigel = 4:1) were injected into the left lung of 4-week-old male C57BL/6 mice. The mice were sacrificed nine days after the injection, and their lungs were obtained for determining if lung cancer was developed.

### Development of a mouse model of traumatic brain injury

Adult male C57BL/6 mice at the weight between 25 - 30 g were anesthetized using 3% tribromoethanol in PBS, 200 μL per 20 g weight, and immobilized by placing the head in a stereotactic frame (Stoelting, Wood Dale, IL). A heating pad was placed under the mouse to maintain body temperature at 37°C. The TBI model was established using a controlled cortical impact (CCI) device (PinPoint Precision Cortical Impactor PCI3000; Hatteras Instruments, Cary, NC, USA). A 3-mm diameter rounded steel impactor tip of a controlled cortical impact (CCI) device (PinPoint Precision Cortical Impactor PCI3000; Hatteras Instruments Inc., Cary, NC, USA) was placed on the exposed intact dura. The cortical surface was hit vertically at an impact velocity of 1.5 m/s, deformation depth of 1.5 mm, and dwell time of 100 ms. Bleeding of the injured cortical surface was controlled by applying a sterile cotton gauze and pressure. The cranial defect was sealed with sterile bone wax, and the incision was closed with interrupted 6-0 silk sutures. The animals were placed in heated cages until they regained full consciousness and then moved to their home cages. The same procedure was performed in sham animals without the CCI injury.

### Administration of LPS and BSO in mice

Male C57BL/6Slac mice at the weight between 18 - 25 g were administered with 0.5 or 1 mg/kg LPS with intraperitoneal (i.p.) injection. The stock solution of LPS with the final concentration of 0.2 mg/ml was made by dissolving LPS in PBS, which were prepared when needed. The mice were administered with these doses of LPS every 24 h for three days.

For administration of BSO in mice, male C57BL/6Slac mice at the weight between 17 - 22 g were administered with 600 or 800 mg/kg BSO with intraperitoneal (i.p.) injection. The stock solution of BSO with the final concentration of 0.2 mg/ml was made by dissolving BSO in PBS. The mice were administered with BSO 12 h and 24 h after the first BSO administration. AF determinations were conducted 1 h after the final BSO administration.

### Exposures of UVC radiation

After C57BL/6 mice, ICR mice, or BALB/cASlac-nu nude mice at the weight between 18 - 35 g were anesthetized briefly with chloral hydrate at the dosage of 1 ml 3.5% (w/v)/100 g, the ears of the mice were exposed to a UVC lamp (TUV 25W /G25 T8, Philips, Hamburg, Germany) with the power density of 0.55±0.05 mW/cm^2^.

### Imaging of the AF of mouse’s skin

The skin’s AF of mouse’s ears was imaged under a two-photon fluorescence microscope (A1 plus, Nikon Instech Co., Ltd., Tokyo, Japan), with the excitation wavelength of 488 nm and the emission wavelength of 500 - 530 nm. The AF was quantified by the following approach: Sixteen spots with the size of approximately 100 × 100 μm^2^ on the scanned images were selected randomly. After the average AF intensities of each layer were calculated, the sum of the average AF intensities of all layers of each spot was calculated, which is defined as the AF intensity of each spot. If the value of average AF intensity of certain layer is below 50, the AF signal of the layer was deemed background noise, which was not counted into the sum. The AF spectra of the mice were determined by using a two-photon fluorescence microscope (A1 plus, Nikon Instech Co., Ltd., Tokyo, Japan). After the imaging, the images were analyzed by the software of the microscope.

### Determinations of the AF and the AF spectrum of human subjects’ skin

A portable AF imaging equipment was used to determine the skin’AF of the human subjects. The excitation wavelength was 485 nm, and the emission wavelength was 500 - 550 nm. By using a portable equipment for measuring AF spectrum, we determined the spectra of the skin’ AF of the human subjects. The emitting light of the wavelength longer than 450 nm was detected, when the excitation wavelength of 445 nm was used.

### Determinations of the serum levels of cytokines of mice

Cytokine concentrations in the serum of mice were determined using a Bio-Plex Pro assay kit (Bio-Rad Laboratories). A total of 50 μl of antibody-conjugated beads was added to the assay plate. After the samples were diluted 1:4, 50 μl of diluted samples, standards, the blank, and the controls was added to the plate. The plate was incubated in the dark at room temperature (RT) with the shaking speed at 300 rpm for 30 min. After three washes with 100 μl wash buffer, a total of 25 μl biotinylated antibody was added to the plate. The plate was incubated in the dark at RT with shaking speed at 300 rpm for 30 min, which was washed 3 times with 100 μl washing buffer. After a total of 50 μl streptavidin-phycoerythrin (PE) was added to the plate, the plate was incubated in the dark at RT with shaking speed at 300 rpm for 10 min. After three washes, the samples were assessed by a Bio-Plex protein array reader. Bio-Plex Manager 6.0 software was used for the data acquisition and analysis of the assay.

### Western blot assays

The lysates of the skin were centrifuged at 12,000 *g* for 20 min at 4°C. The protein concentrations of the samples were quantified using BCA Protein Assay Kit (Pierce Biotechonology, Rockford, IL, USA). As described previously^22^, a total of 10 μg of total protein was electrophoresed through a 10% SDS-polyacrylamide gel, which were then electrotransferred to 0.45 μm nitrocellulose membranes (Millipore, CA, USA). The blots were incubated with a monoclonal Anti-Cytokeratin 1 (ab185628, Abcam, Cambridge, UK) (1:4000 dilution) or actin (1:2000, sc-58673, Santa Cruz Biotechnology, Inc., Dallas, TX, USA) with 0.05% BSA overnight at 4°C, then incubated with HRP conjugated Goat Anti-Rabbit IgG (H+L) (1:4000, Jackson ImmunoResearch, PA, USA) or HRP conjugated Goat Anti-mouse IgG (1:2000, HA1006, HuaBio, Zhejiang Province, China). An ECL detection system (Thermo Scientific, Pierce, IL, USA) was used to detect the protein signals. The intensities of the bands were quantified by densitometry using Image J.

### Immunohistochemistry

Ten μm paraffin-sections of skin were obtained by a Leica Cryostat, mounted onto poly-L-lysine coated slides and stored at room temperature. After the sections were incubated in Xylene three times, the skin sections were dehydrated sequentially in 100% EtOH, 95% EtOH and 70% EtOH. After two washes with PBS, the sections were blocked in 10% goat serum for 1 hr, which were incubated in monoclonal Anti-Cytokeratin 1 (ab185628, abcam, Cambridge, UK) (1:1000 dilution), containing 1% goat serum at 4 °C overnight. After three washes in PBS, the sections were incubated with Alexa Fluor 647 goat anti-rabbit lgG (1:1000 dilution) (Invitrogen, CA,USA) for 1 hr in darkness at RT, followed by staining in 0.2% DAPI solution (Beyotime, Haimen, Jiangsu Province, China) for 5 min. The sections were mounted in Fluorescence Mounting Medium (Beyotime, Haimen, Jiangsu Province, China). To compare the intensity of fluorescence of each sample, at least three randomly picked fields in each section were photographed under a Leica microscope.

### Histology

Skin biopsies from the ears of the mice were obtained, which were placed immediately in 4% (w/v) paraformaldehyde buffer. After 12 - 24 h, paraffin embedding procedure was conducted on the samples. Hematoxylin / Eosin staining was performed according to the manufacturer’s protocol (Beyotime, Haimen, Jiangsu Province, China).

### Laser-based delivery of keratin 1 siRNA into mouse’s skin

Male C57BL/6 mice were briefly anesthetized with 3.5% (w/v) chloral hydrate (0.2 ml / 20 g). After exposure to laser, the ears of the mouse were transfected with K1 siRNA using Lipofectamine 3000 following the manufacturer’s instructions (Thermo Fisher Scientific, Waltham, MA, USA). The sequences of the mouse K1 siRNA were CUCCCAUUUGGUUUGUAGCTT and UGACUGGUCACUCUUCAGCTT (GenePharma, Shanghai, China).

### Statistical analyses

All data are presented as mean ± SEM. The data from animal and cell culture studies were assessed by one-way ANOVA, followed by Student - Newman - Keuls *post hoc* test, except where noted. *P* values less than 0.05 were considered statistically significant. The data from the studies on human subjects were assessed by Kruskal-Wallis multiple comparison test (K-W test) or Mann-Whitney test. *P* values less than 0.05 were considered statistically significant.

## Results

### 1 The epidermal green AF intensity was significantly associated with both LPS dosages and serum levels of multiple cytokines in LPS-administered C57BL/6 mice

We determined the effects of LPS, an inducer of inflammation, on the green AF of the skin of C57BL/6 mice’s ears: LPS produced dose-dependent increases in the AF intensity 3 d after the LPS administration (**Figs. 1A and 1B)**, with the AF intensity being significantly associated with the LPS dosages (**Fig. 1C)**. The spatial distribution of the LPS-induced AF is distinctly polyhedral, exhibiting the characteristic structure of the suprabasal keratinocytes at the stratum spinosum (**Fig. 1A)**. Both the LPS dosages and the green AF intensity were also significantly associated with the Log-transformed serum levels of multiple cytokines of the mice, including IL-1β, IL-2, IL-5, IL-6, IL-10, IL-12(p40), monocyte chemoattractant protein-1 (MCP-1) and G-CSF, assessed at 3 d after the LPS administration (**Table 1**). Moreover, both the LPS dosages and the green AF intensity were significantly associated with the serum levels of MIP-1α, MIP-1β and RANTES/CCL5 (**Table 2**). In contrast, neither the LPS dosage nor the epidermal AF intensity was significantly associated with either the serum levels or the Log-transformed serum levels of certain cytokines 3 d after the LPS administration (**Supplementary Tables 1 and 2**).

**Table 1.** Epidermal green AF intensity was significantly associated with the Log-transformed serum levels of multiple cytokines in LPS-administered C57BL/6 mice. The epidermal green AF intensity was significantly associated with the Log-transformed serum levels of IL-1β, IL-2, IL-5, IL-6, IL-10, IL-12(p40), monocyte chemoattractant protein-1 (MCP-1) and G-CSF of C57BL/6 mice’s. Three d after the mice were i.p. administered with 0.5 or 1 mg/kg LPS, the epidermal green AF intensity of the ear’s skin of the mice was determined. The serum levels of multiple cytokines of the mice were also determined. N = 4-6.

| Cytokine | [LPS] vs. Log [Cytokine] |  | AF vs. Log [Cytokine] |  |
| --- | --- | --- | --- | --- |
|  | *p value* | R | *p value* | R |
| IL-1β | 0.0007 | 0.7725 | 0.0141 | 0.6181 |
| IL-2 | 0.0006 | 0.7815 | 0.0153 | 0.6122 |
| IL-5 | 0.0143 | 0.6169 | 0.0258 | 0.5723 |
| IL-6 | <0.0001 | 0.8608 | 0.0427 | 0.5289 |
| IL-10 | 0.0029 | 0.7122 | 0.0081 | 0.6549 |
| IL-12(p40) | 0.0043 | 0.6910 | 0.0106 | 0.6374 |
| G-CSF | <0.0001 | 0.9250 | 0.0183 | 0.5990 |
| MCP-1 | 0.0040 | 0.6949 | 0.0342 | 0.5485 |

**Table 2.** Epidermal green AF intensity was significantly associated with the serum levels of three cytokines in LPS-administered C57BL/6 mice. Epidermal green AF intensity was significantly associated with the serum levels of MIP-1α, MIP-1β and RANTES/CCL5. Three d after the mice were i.p. administered with 0.5 or 1 mg/kg LPS, the epidermal green AF intensity of the ear’s skin of the mice was determined. The serum levels of the cytokines of the mice were also determined. N = 4-6.

| Cytokine | [LPS] vs. [Cytokine] |  | AF vs. [Cytokine] |  |
| --- | --- | --- | --- | --- |
|  | *p value* | R | *p value* | R |
| MIP-1α | 0.0032 | 0.7072 | 0.0012 | 0.7542 |
| MIP-1β | 0.0080 | 0.6555 | 0.0031 | 0.7085 |
| RANTES | 0.0003 | 0.8075 | 0.0002 | 0.8184 |

**Figure 1.**
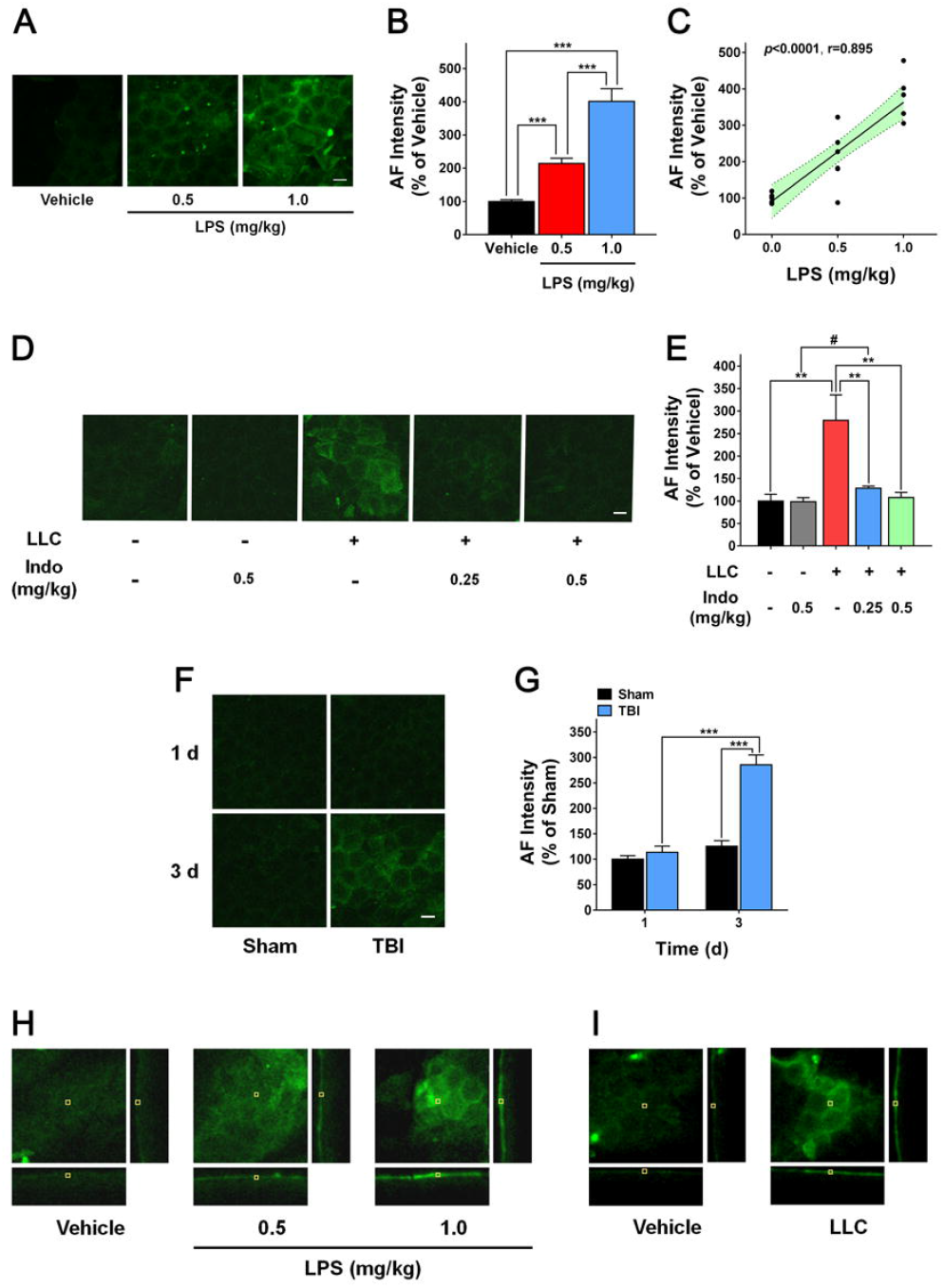
The green AF intensity of the skin was significantly associated with both LPS dosages and serum levels of multiple cytokines in LPS-administered C57BL/6 mice. (A) LPS produced increased the green AF of the mice’s ears 3 d after i.p. injections of 0.5 or 1 mg/kg LPS. Scale bar = 20 μm. (B) LPS produced significant and dose-dependent increases in the epidermal green AF intensity of the mice’s ears 3 d after the LPS administration. N = 16-19. ***, *P* < 0.001. (C) The AF intensity was significantly associated with the LPS dosages. The data were obtained from one representative experiment out of the three independent experiments. N = 6. The *P* values of the other two independent experiments were < 0.001. The r values of the other two independent experiments were 0.919 和 0.816, respectively. (D, E) There were marked increases in the green AF of the ears’ skin of all of the mice that had developed lung cancer, which were abolished by administration of the anti-inflammation drug indomethacin (Indo), assessed at 8 d after the LLC injection. Indo was administered 3 d after the LLC injection. N = 4 - 7. #, *P* < 0.05 (Student t-test); **, *P* < 0.01. Scale bar = 20 μm. (F, G) Traumatic brain injury of mice led to a significant increase in the green AF of ears’ skin of the mice, assessed at 3 d after head injury. N = 6 - 9. ***, *P* < 0.001. Scale bar = 20 μm. (H,I) Orthographic green AF images of the skin of the mice’s ears showed that the increased AF induced by LPS or lung cancer occurred only at the locations that were approximately 10 - 20 μm apart from the outer surface of the epidermis, assessed at 1 d after LPS administration or 9 d after LLC injections. The images of XY axis (150 μm X 150 μm), YZ axis (150 μm X 100 μm) and XZ axis (150 μm X 100 μm) were shown. N = 6.

Inflammation plays important roles in both pathogenesis and development of lung cancer 23,24. We found that development of lung cancer in mice (**Supplementary Fig. 1A**) also produced a significant increase in the skin’s green AF with polyhedral structure (**Fig. 1D and Fig. 1E**). Administration of the anti-inflammation drug indomethacin virtually abolished the lung cancer-induced AF increase (**Fig. 1D and Fig. 1E**), while the drug did not affect the weight of the lung with lung cancer (**Supplementary Fig. 1B**), thus indicating a key role of inflammation in the lung cancer-induced AF increase. Traumatic brain injury, a brain disorder with markedly increased inflammatory responses^25^, also led to a significant increase in the green AF with polyhedral structure of the mice’s skin 3 d after head injury (**Fig. 1F and Fig. 1G**).

We further investigated the localization of the increased AF induced by LPS and lung cancer in the mouse models: Orthographic green AF images of the mice’s skin showed that both LPS- and lung cancer-induced AF occurred only at certain layer of the skin with the thickness of approximately 10 μm, which was approximately 10 - 20 μm apart from the outer surface of the epidermis (**Figs. 1F and 1G**). H&E staining of the skin showed that the thickness of the epidermis was approximately 25 - 30 μm, while the thickness of the stratum corneum was less that 3 μm (**Supplementary Fig. 1C**), indicating that the origin of the AF was localized at the epidermis between the stratum corneum and the dermis.

### 2. Green AF intensity of mouse’s skin or skin cell cultures was highly associated with the dosages of oxidative stress inducers

We determined the effects of oxidative stress on the green AF of B16-F10 cells, a skin cell line: H2O2 produced concentration-dependent increases in the green AF intensity (**Figs. 2A and Fig. 2B)**, with the AF intensity being highly associated with the H2O2 concentrations both 1 and 3 h after the H2O2 exposures (**Figs. 2C)**. UVC irradiation, an oxidative stress inducer of the skin^26,27^, also produced dose-dependent increases in the green AF intensity of the cells (**Supplementary Figs. 2A and 2B)**, with the AF intensity being highly associated with the UVC dosages 0.3, 1 and 6 h after the UVC irradiation (**Supplementary Fig. 2C**).

**Fig 2.**
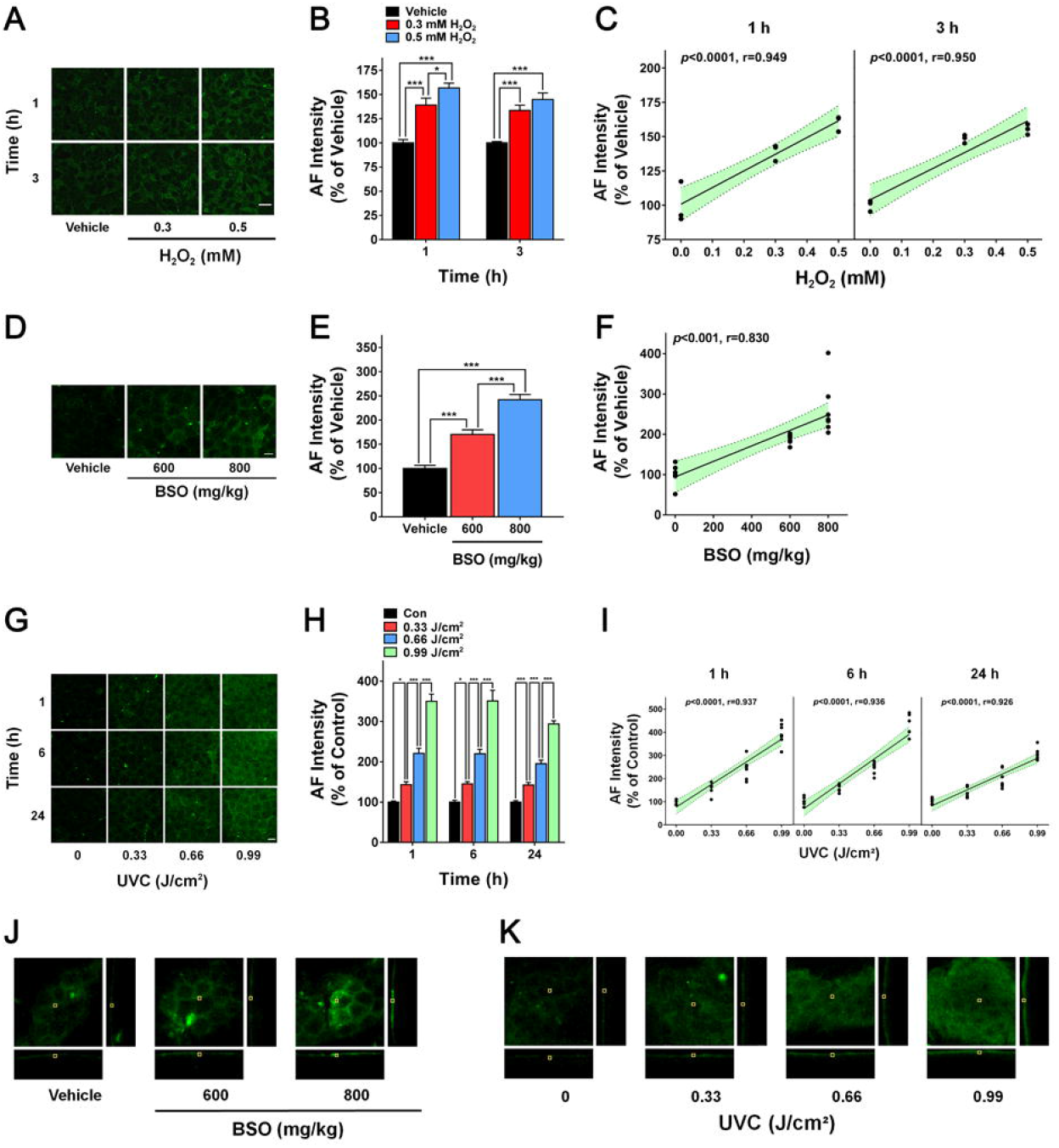
Green AF intensity of mouse’s skin or skin cell cultures was highly associated with the dosages of oxidative stress inducers. (A) H2O2 produced increased green AF intensity of B16-F10 cells 1 and 3 h after the H2O2 exposures. Scale bar = 20 μm. (B) H2O2 produced significant and concentration-dependent increases in the green AF intensity of B16-F10 cells 1 and 3 h after the H2O2 exposures. N = 9. *, *P* < 0.05; ***, *P* < 0.001. The data were collected from three independent experiments. (C) The AF intensity was significantly associated with the H2O2 concentrations 1 and 3 h after the H2O2 exposures. The data were obtained from one representative experiment out of the three independent experiments. N = 3. The *P* values of the other two independent experiments were < 0.001. The r values of the other two independent experiments conducted 1 h after the H2O2 exposures were 0.923, respectively; and r values of the other two independent experiments conducted 3 h after the H2O2 exposures were 0.963 and 0.942, respectively. (D,E) Intraperitoneal injection of BSO produced dose-dependent increases in the epidermal green AF of mice’s ears. N = 19-21. ***, *P* < 0.001. Scale bar = 20 μm. (F) The intensity of the epidermal green AF was significantly associated with the BSO dosages. The data were obtained from one representative experiment out of three independent experiments. N = 6-7. The *P* values of the other two independent experiments were < 0.001. The r values of the other two independent experiments were 0.733 and 0.850, respectively. (G) UVC irradiation produced increased green AF intensity of the ear’s skin of C57BL/6 mice 1, 6, or 24 h after the UVC exposures. Scale bar = 20 μm. (H) UVC irradiation produced significant and dose-dependent increases in the green AF intensity of the ear’s skin of C57BL/6 mice 1, 6, or 24 h after the UVC exposures. N = 10 - 13. *, *P* < 0.05; ***, *P* < 0.001. (I) The AF intensity was significantly associated with the UVC dosages 1, 6, or 24 h after the UVC exposures. The data were obtained from one representative experiment out of two independent experiments. N = 5-7. The *P* values of the other independent experiments were < 0.001. The r values at 1 h, 6 h, and 24 h were 0.930, 0.961 and 0.896 respectively. (J,K) Orthographic green AF images of the mice’s skin showed that the increased AF induced by BSO UVC occurred only at the locations that were approximately 10 - 20 μm apart from the outer surface of the epidermis, assessed at 1 d after BSO administration or 6 h after UVC irradiation. The images of XY axis (150 μm X 150 μm), YZ axis (150 μm X 100 μm) and XZ axis (150 μm X 100 μm) were shown. N = 6.

Intraperitoneal injection of L-(S,R)-Buthionine Sulfoximine (BSO), an irreversible glutathione synthesis inhibitor, produced dose-dependent increases in the green AF intensity of the ear’s skin of C57BL/6 mice (**Figs. 2D and 2E**), with the AF intensity being significantly associated with the BSO dosages (**Fig. 2F**). UVC also produced dose-dependent increases in the green AF intensity of the ear’s skin (**Figs. 2G and 2H**), with its dosages being highly associated with the AF intensity 1, 6 and 24 h after the UVC exposures (**Fig. 2I**).

The spectra of both basal and UVC-induced AF of C57BL/6 mice, ICR mice and nude mice were similar, reaching maximal AF intensity at 470 - 500 nm when the excitation wavelength was 800 nm under a two-photon fluorescence microscope (**Supplementary Fig. 2D**). Orthographic green AF images of the mouse’s skin showed that both BSO- and UVC-induced AF occurred only at certain layer of the skin (**Figs. 2J and 2K)**, which were highly similar with those of the LPS- and lung cancer-induced AF images, indicating that the origin of the AF was also localized at the epidermis between the stratum corneum and the dermis. This observation, combined with our finding that the spatial distribution of the AF induced by LPS, BSO, UVC, or lung cancer exhibited the characteristic structure of the suprabasal keratinocytes at the stratum spinosum (**Figs. 1A, 1D, and 1F and Figs. 2D and 2G)**, has indicated that the AF is localized at the stratum spinosum.

### 3. Marked and asymmetrical increases in the green AF intensity of the Dorsal Index Fingers’ skin of AIS patients hold promise as a novel biomarker for AIS diagnosis

There are significant increases in inflammatory responses and oxidative stress in the body of AIS patients^28-30^ and lung cancer patients^14,23^,24. To determine if our findings from the animal and cell culture studies are also applicable to major inflammation- and oxidative stress-associated diseases, we determined if there is also increased green AF in the skin of AIS patients and lung cancer patients: In the skin of at least one of their Dorsal Index Fingers, the percentage of the persons with the polyhedral structure of strong AF intensity in the AIS Group was remarkably higher than that of the Lung Cancer Group, the High-Risk Group, the Recovery Group, or the Healthy Group (**Fig. 3A**). In contrast, compared with the other four groups, there was a higher percentage of the lung cancer patients who had polyhedral structure with moderate AF intensity (**Fig. 3A**).

**Fig 3.**
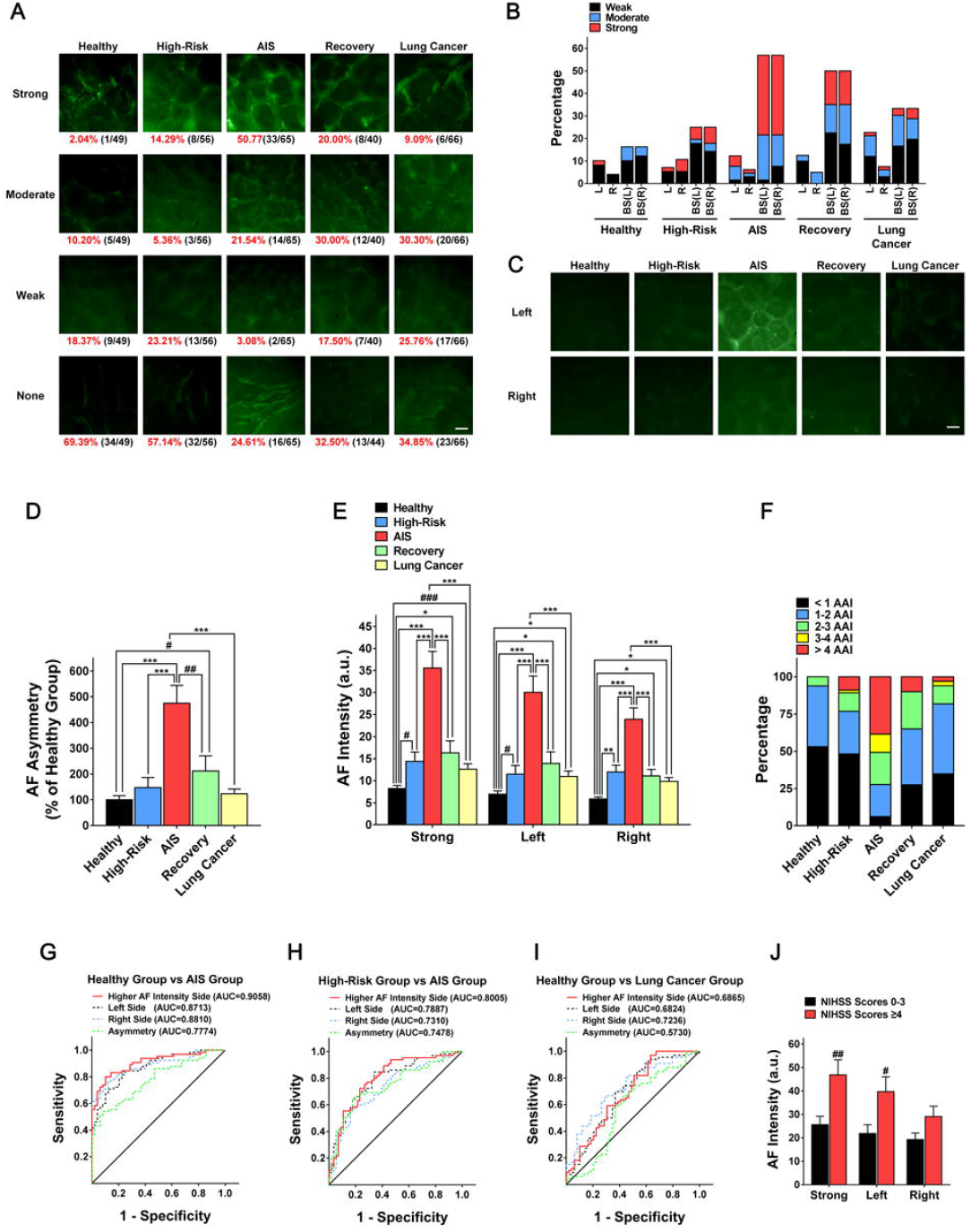
Profound and asymmetrical increases in the green AF intensity of the Dorsal Index Fingers’ skin of AIS patients hold promise as a novel biomarker for AIS diagnosis. (A) Among the Healthy Group, the High-Risk Group, the AIS Group, the Recovery Group and the Lung Cancer Group, there were marked differences in the percentages of the AF without polyhedral structure, the AF with polyhedral structure of ‘weak AF intensity’ - the AF intensity that was 100 – 150 % of the average AF intensity of the right and left Dorsal Index Fingers’ skin of the healthy controls, the AF with polyhedral structure of ‘moderate AF intensity’ - the AF intensity that was at 151 – 300 % of the average AF intensity of the right and left Dorsal Index Fingers’ skin of the healthy controls., and the AF with polyhedral structure of ‘strong AF intensity’ - the AF intensity that was at least 300% of the average AF intensity of the right and left Dorsal Index Fingers’ skin of the healthy controls. Scale bar = 100 μm. (B) In these groups, the distributions of the polyhedral structure with highly, moderately or weakly increased AF in their right and left hands were differential. The number of the subjects in the Healthy Group, the High-Risk Group, the AIS Group, the Recovery Group, and the Lung Cancer Group was 49, 56, 65, 40 and 66, respectively. (C) Representative green AF images of the left and right Dorsal Index Fingers of the Healthy Group, the High-Risk Group, the AIS Group, the Recovery Group, and the Lung Cancer Group. N = 49-66. (D) The AF asymmetry of the Dorsal Index Fingers’ skin of the AIS patients was markedly higher than that of the other four groups. N = 49-66. ***, *P* < 0.001 (K-W test); #, *P* < 0.05 (Mann-Whitney test); ##, *P* < 0.01 (Mann-Whitney test). (E) The AF intensity of their Stronger AF Side of the AIS patients was significantly higher than that of the other four groups. N = 49-66. #, *P* < 0.05 (Mann-Whitney test); ###, *P* < 0.001 (Mann-Whitney test); *, *P* < 0.05 (K-W test); **, *P* < 0.01 (K-W test); ***, *P* < 0.001 (K-W test). (F) In comparisons of the AF intensity of their Stronger AF Sides, a remarkably higher percentage of the AIS patients had their AF intensity higher than 200% of the average AF intensity of the healthy controls (AAI) in the five groups of subjects. N = 49-66. (G) ROC analysis using the AF intensity of the Stronger AF Side, the AF intensity of the left Dorsal Index Fingers’ skin, the AF intensity of the right Dorsal Index Fingers’ skin, or the AF asymmetry as the sole parameter showed that the AUC was 0.9058, 0.8713, 0.8810, and 0.7774, respectively, for differentiating the healthy controls and the AIS patients. N =49-65. (H) ROC analysis using the AF intensity of the Stronger AF Side, the AF intensity of the left Dorsal Index Fingers’ skin, the AF intensity of the right Dorsal Index Fingers’ skin, or the AF asymmetry as the sole parameter showed that the AUC was 0.8006, 0.7887, 0.7310 and 0.7478, respectively, for differentiating the High-Risk subjects and the AIS patients. N = 56-65. (I) ROC analysis using the AF intensity of the Stronger AF Side, the AF intensity of the left Dorsal Index Fingers’ skin, the AF intensity of the right Dorsal Index Fingers’ skin, or the AF asymmetry as the sole parameter showed that the AUC was 0.6865, 0.6824, 0.7236, and 0.5730, respectively, for differentiating the healthy controls and the lung cancer patients. (J) Compared with the group of the AIS patients with the NIHSS scores between 0 - 3, the group of the AIS patients with the NIHSS scores equal to or higher than 4 had significantly higher green AF intensity of both their Stronger AF Side and their left Dorsal Index Fingers. N =31-32. #, *P* < 0.05 (Mann-Whitney test); ##, *P* < 0.01 (Mann-Whitney test).

Among these five groups, the distributions of the polyhedral structure with strong, moderate or weak AF intensity in their right and left Dorsal Index Fingers were differential (**Fig. 3B**). The AF intensity in their right and left Dorsal Index Fingers of a majority of the AIS patients was distinctly different, as shown in the representative images (**Fig. 3C**), which led us to define the difference (in absolute value) between the AF intensity of the right and left side of the skin of a subject as ‘AF Asymmetry’, with the side of the skin with higher AF intensity being defined as ‘Stronger AF Side’. We found that the AF Asymmetry of the Dorsal Index Fingers’ skin of the AIS patients was approximately 263 - 352% higher than that of the other four groups (**Fig. 3D**).

The AF intensity of the Stronger AF Side of the AIS patients was significantly higher than that of the other four groups (**Fig. 3E**). However, we did not observe any correlation between the side with higher AF intensity and either the side of the AIS patient’s brain with blood clots, as assessed by MRI, or the opposite side (data not shown). The AF intensity of both right and left Dorsal Index Fingers’ skin of the AIS patients was also significantly higher than that of the other four groups (**Fig. 3E**). Moreover, the green AF intensity of both right and left Dorsal Index Fingers’ skin of the lung cancer patients was significantly higher than that of the healthy controls (**Fig. 3E**). Among the five groups of subjects, remarkably higher percentages of the AIS patients had their AF intensity on their Stronger AF Side that was at least 200% higher than the average AF intensity of the healthy controls (**Fig. 3F**).

ROC analyses using the AF intensity of the Stronger AF Side, the AF intensity of the left Dorsal Index Fingers’ skin, the AF intensity of the right Dorsal Index Fingers’ skin, or the AF asymmetry as the sole diagnostic parameter showed the following Area Under Curve (AUC) values: The AUC values were 0.9058, 0.8713, 0.8810, and 0.7774, respectively, for differentiating the healthy controls and the AIS patients (**Fig. 3G**); the AUC were 0.8005, 0.7887, 0.7310 and 0.7478, respectively, for differentiating the High-Risk subjects and the AIS patients (**Fig. 3H**); and the AUC values were 0.6865, 0.6824, 0.7236, and 0.5730, respectively, for differentiating the healthy controls and the lung cancer patients (**Fig. 3I**). The sensitivity and specificity for differentiating the AIS patients and the healthy controls were 0.8308 and 0.8571, respectively (**Table 3**).

**Table 3.** The sensitivity and specificity for differentiating the AIS patients and the healthy controls, differentiating the AIS patients and the High-Risk Group, and differentiating the lung cancer patients and the healthy controls. The sensitivity and specificity for differentiating the AIS patients and the healthy controls were 0.8308 and 0.8571, respectively.

|  | **Cutoff >** | **Sensitivity** | **95% CI** | **Specificity** | **95% CI** |
| --- | --- | --- | --- | --- | --- |
| **Healthy Group vs AIS Group** |  |  |  |  |  |
| Higher AF Intensity Side | > 13.1 | 0.8308 | 0.7173 to 0.9124 | 0.8571 | 0.7276 to 0.9406 |
| Left Side | > 10.8 | 0.7846 | 0.6651 to 0.8769 | 0.7959 | 0.6566 to 0.8976 |
| Right Side | > 8.513 | 0.8154 | 0.6997 to 0.9008 | 0.8163 | 0.6798 to 0.9124 |
| Asymmetry | > 4.415 | 0.6923 | 0.5655 to 0.8009 | 0.6735 | 0.5246 to 0.8005 |
| **High-Risk Group vs AIS Group** |  |  |  |  |  |
| Higher AF Intensity Side | > 15.27 | 0.7538 | 0.6313 to 0.8523 | 0.7321 | 0.597 to 0.8417 |
| Left Side | > 11.98 | 0.7231 | 0.5981 to 0.8269 | 0.7143 | 0.5779 to 0.827 |
| Right Side | > 12.22 | 0.6769 | 0.5495 to 0.7877 | 0.6607 | 0.5219 to 0.7819 |
| Asymmetry | > 4.605 | 0.6923 | 0.5655 to 0.8009 | 0.7321 | 0.597 to 0.8417 |
| **Healthy Group vs Lung Cancer Group** |  |  |  |  |  |
| Higher AF Intensity Side | > 9.142 | 0.6212 | 0.4934 to 0.7378 | 0.6122 | 0.4624 to 0.748 |
| Left Side | > 6.326 | 0.697 | 0.5715 to 0.8041 | 0.6327 | 0.4829 to 0.7658 |
| Right Side | > 7.131 | 0.6667 | 0.5399 to 0.778 | 0.6939 | 0.5458 to 0.8175 |
| Asymmetry | > 2.219 | 0.6061 | 0.4781 to 0.7242 | 0.5918 | 0.4421 to 0.73 |

We further determined if the AF intensity was associated with the patients’ scores of National Institutes of Health Stroke Scale (NIHSS) - a scale for rating stroke severity^31^. Compared with the group of the AIS patients with the NIHSS scores between 0 - 3, the group of the AIS patients with the NIHSS scores equal to or higher than 4 had significantly higher green AF intensity of their Stronger AF Side and their left Dorsal Index Fingers (**Fig. 3J**). The spectrum of the AF of the AIS patient’s skin was virtually identical as that of the healthy controls (**Supplementary Fig. 3**).

### 4. K1 is an origin of the UVC-induced increase in the epidermal AF of mice

Among the four known epidermal autofluorescent molecules^10^, our study excluded the possibility that melanin is the origin of the UVC-induced green AF of mice, since UVC was capable of increasing the AF of the ear’s skin of ICR mice that are melanin-deficient^36^ (**Figs. 4A and 4B)**. Our observations further excluded the possibility that FAD is the origin of the UVC-induced green AF: UVC significantly decreased the FAD level of the skin (**Fig. 4C**), when it significantly increased the epidermal AF intensity of C57BL/6 mouse’s ears 1 h after the UVC exposures (**Figs. 2G and 2H**). NADPH is unlikely the origin of the AF, since the excitation wavelength for NAD(P)H’ AF is far below the excitation wavelength used in our study^35^.

**Fig 4.**
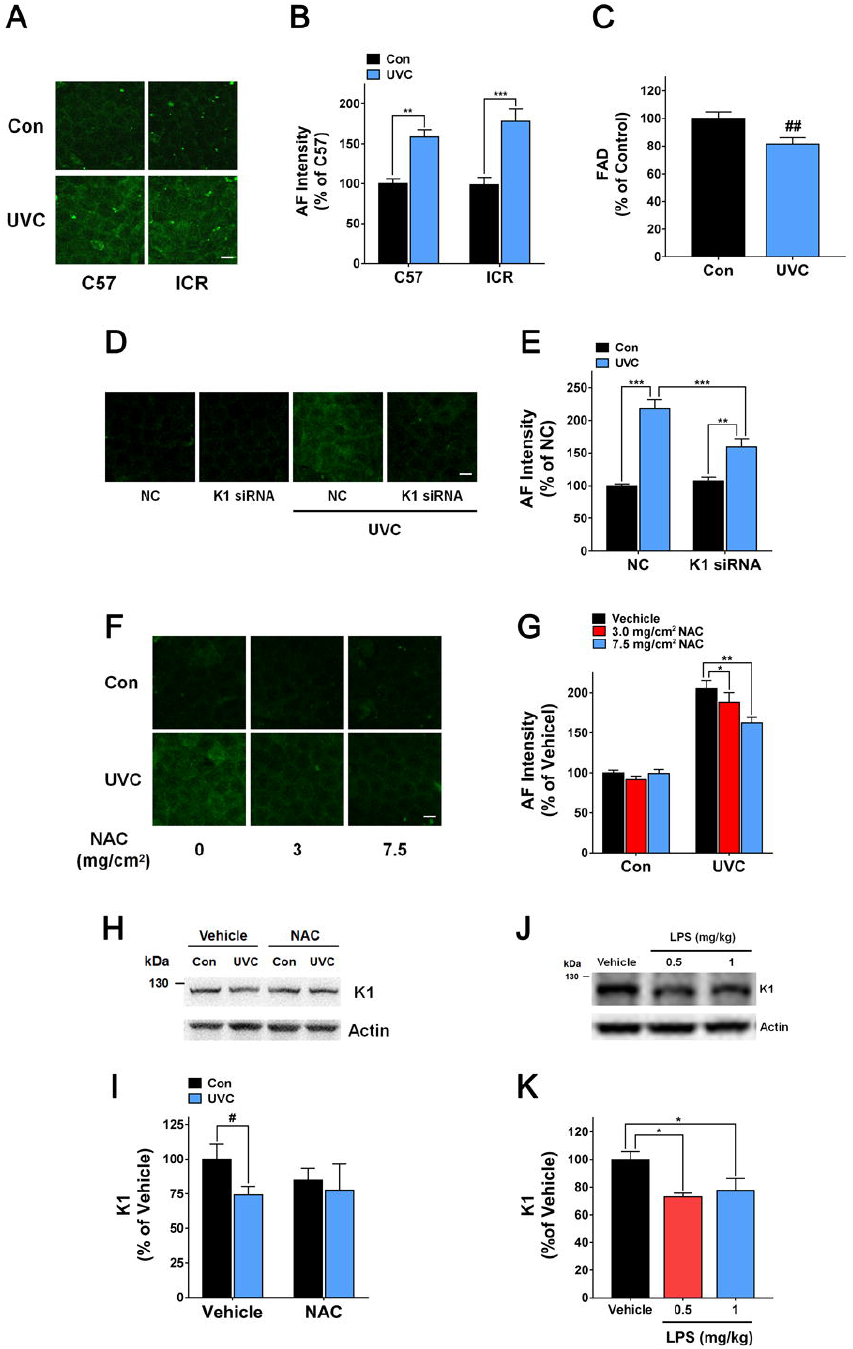
K1 is an origin of the UVC-induced increase in the epidermal AF of C57BL/6 mice. (A, B) UVC produced increased green AF intensity of the skin of ICR mice 3 - 6 h after the UVC exposures. N = 4 - 5. **, *P* < 0.01; ***, *P* < 0.001. Scale bar = 20 μm. (C) UVC significantly decreased the FAD level of the skin, assessed at 1 h after the UVC irradiation. N = 21. ##, *P* < 0.01 (Student t-test). (D, E) K1 siRNA administration significantly attenuated the UVC-induced AF increase. N = 6 -10. **, *P* < 0.01; ***, *P* < 0.001. (F, G) NAC administration dose-dependently attenuated the UVC-induced AF increase. N = 6; *, *P* < 0.05; **, *P* < 0.01. (H, I) NAC administration significantly and dose-dependently attenuated the UVC-induced K1 decrease. N = 8; #, *P* < 0.05 (Student t-test). (J, K) LPS induced a significant decrease in the K1 levels of the ears 1 d after the mice were i.p. administered with 0.5 or 1 mg/kg LPS. One d after the mice were i.p. administered with 0.5 or 1 mg/kg LPS, the mice were i.p. administered with 0.5 or 1 mg/kg LPS again. One h after the administration, the K1 levels of the ears were determined by Western blot assay. N = 6. *, *P* < 0.05.

Several lines of evidence has suggested that K1 and/or K10 may be the origin of the oxidative stress- and inflammation-induced increases in epidermal AF: 1) The AF is selectively localized at the stratum spinosum (**Fig. 1A and Figs. 2D and 2G)**, where the only autofluorescent molecules that are selectively localized at the stratum spinosum are K1 and K10^21^; 2) the spectrum of the UVC-induced green AF (**Supplementary Fig. 2F**) matches that of keratins’ AF^32^; and 3) the structure and localization of the LPS- (**Fig. 1A**), BSO- (**Fig. 2G**) and UVC-induced AF (**Fig. 2G**) match the structure and localization of K1/K10 heterodimers in the epidermis, which form unique, highly dense bundles that are parallel with the cell membranes of the polyhedral suprabasal cells^33,34^. Immunostaining assays of the ear’s skin of C57BL/6 mice using anti-K1 antibody also showed that the polyhedral structure of the K1-positive regions (**Supplemental Fig. 4A**) matched the polyhedral structure of the LPS- (**Fig. 1A**), BSO- (**Fig. 2G**) and UVC-induced AF (**Fig. 2G**).

Since K1 is associated with K10 as K1/K10 heterodimers in differentiated keratinocytes^21,37,38^, we determined if K1 and K10 exist as monomers or heterodimers in the ear’s skin of C57BL/6 mice. Western blot assays on K1 using normal denaturing conditions showed that in the samples of the ear’s skin, there was a distinct band at approximately 127 kDa, while the level of K1 monomers at the 67 kDa band was exceedingly low (**Supplementary Figs. 4A)**. When the samples were prepared in 8 M urea, Western blot assays on K1 showed that the 127 kDa band disappeared with appearance of both a 67 kDa and a 59 kDa band, while Western blot assays on K10 showed an increased band at 60 kDa (**Supplementary Figs. 4B)**. These observations have indicated that the 127 kDa band is the band of the K1/K10 heterodimers.

To determine the role of K1 in the UVC-induced AF increase, we applied a laser-based technology to deliver K1 siRNA into the skin of the mouse’s ears. Both Western blot (**Supplementary Figs. 4C and 4D)** and immunostaining assays (**Supplementary Fig. 4E**) showed that the laser-based K1 siRNA administration led to a K1 decrease. The administration of the K1 siRNA also significantly attenuated the UVC-induced AF increase (**Figs. 4D and 4E**). We further determined the effects of the antioxidant N-acetyl cysteine (NAC) on the UVC-induced AF: NAC administration dose-dependently attenuated the UVC-induced AF increase (**Figs. 4F and 4G**). Surprisingly, UVC induced a significant decrease in the K1 level of the ears, which was blocked by the NAC administration (**Figs. 4H and 4I**). LPS also induced a significant decrease in the K1 level of the ears 1 d after the mice were i.p. administered with 0.5 or 1 mg/kg LPS (**Figs. 4J and 4K**). Both H2O2 (**Supplementary Figs. 4H and 4I)** and UVC (**Supplementary Figs. 4J and 4K)** induced significant K1 decreases in B16-F10 cells.

## Discussion

This study has provided several lines of evidence indicating the epidermal green AF as a novel endogenous reporter of the levels of inflammation and oxidative stress in the body: First, in skin cell culture studies, oxidative stress produced dose-dependent increases in the green AF, with the AF intensity being highly associated with the dosages of oxidative stress inducers; second, in animal model studies, both inflammation and oxidative stress inducers produced dose-dependent increases in the epidermal green AF, with the AF intensity being significantly associated with the dosages of the inducers; third, both lung cancer and traumatic brain injury induced marked increases in the epidermal green AF of mice; fourth, the lung cancer-induced increase in the epidermal green AF was abolished by an anti-inflammation drug, indicating that inflammation mediates the lung cancer-induced AF increase; fifth, there were significant increases in the green AF of the Dorsal Index Fingers’ skin of both AIS and lung cancer patients, which have shown promising diagnostic value; sixth, the green AF of the Dorsal Index Fingers’ skin of the ischemic stroke patients in recovery phase was significantly decreased, compared with that of the AIS patients; seventh, the green AF of the Dorsal Index Fingers’ skin of the AIS patients is asymmetrical, which is also a novel and promising diagnostic factor; and eighth, the skin’s AF intensity of the High-Risk Group was significantly higher than the Healthy Group, while it was significantly lower than that of the AIS Group, suggesting that we may differentiate non-invasively and efficiently the healthy controls and the High-Risk human subjects by determining their skin’s green AF. Moreover, the increased skin’s AF of the AIS patients and the lung cancer patients had highly similar polyhedral structure as that found in the skin of the LPS- or ROS-exposed mice or the mice with lung cancer or head injury. These findings have collectively suggested that the AF is a common biomarker of the pathological insults in the body of both human and the mice.

Our study has provided first evidence suggesting profound and extensive applications of the keratin’s AF for non-invasive diagnosis of inflammation- and oxidative stress-associated diseases: For differentiating the AIS patients and the healthy controls, the AUC value was 0.9058, when the AF intensity was the sole diagnostic parameter. Notably, the AIS patients and the lung cancer patients were significantly different from each other in both their AF intensity and AF asymmetry, suggesting that the patients of different diseases may have differential properties of their AF. Moreover, our determinations on the green AF of other positions of the skin such as Vetroforefingers have also shown significantly higher green AF in the skin of AIS patients compared with the healthy controls and the High-Risk subjects (data not shown). With increasing information regarding the skin’s green AF at both Dorsal Index Fingers and other positions, analyses of the structural properties of the AF images, additions of the information of clinical symptoms and applications of AI, it is expected that the sensitivity and specificity of this novel diagnostic method may be further enhanced.

Increased inflammation and/or oxidative stress have also been found in the patients’ body of numerous diseases such as Parkinson’s disease^15,16^ and COVID-19^39,40^. Therefore, our AF-based approach may also be used for evaluating the pathological state of the patients of these diseases, which has been supported by our preliminary studies showing that each of the diseases studied has its characteristic changes of the pattern of their skin’s green AF^41,42^. Our study has also indicated that the keratin’s AF-based approach may become the first non-invasive method for monitoring the levels of multiple cytokines in the body, which has further indicated the great potential of this AF imaging technology for non-invasive evaluations of patients’ key biological parameters. Compared with major medical imaging technology including MRI, CT and PET technology, our AF-based imaging technology are more economic, time-efficient and non-invasive.

Our study has found asymmetrical increases in the green AF in the Dorsal Index Fingers of the AIS patients, which is the first report regarding the asymmetrical state of biomedical optical properties of the skin. Sine the AF asymmetry of the AIS patients was remarkably higher than that of the healthy subjects, the High-Risk subjects, and the lung cancer patients, the AF asymmetry has shown significant diagnostic value. The markedly higher AF asymmetry of the AIS patients than that of the High-Risk subjects has also suggested that the AF asymmetry may be mainly originated from ischemic stroke, but not a pre-existing condition. However, we did not find any correlation between the side with higher AF intensity and either the side of the AIS patient’s brain with blood clots or the opposite side. Investigation of the mechanisms underlying the AF asymmetry may expose novel pathological mechanisms underlying AIS.

Our study has indicated K1 as an origin of the oxidative stress- and inflammation-induced epidermal AF increases: First, the K1 siRNA-produced K1 decrease significantly attenuated the UVC-induced AF increase; second, when it attenuated UVC-produced K1 decrease, NAC significantly attenuated the UVC-induced AF increase; third, the increased AF is selectively localized at the stratum spinosum, where the only known autofluorescent molecules that are selectively localized are K1 and K10^21^; fourth, the spectrum of the UVC-induced green AF matches that of keratins’ AF^32^, and fifth, the structure and localization of the LPS- and UVC-induced AF match those of K1/K10 heterodimers in the epidermis^33,34^. Moreover, our studies have excluded the possibility that the other three known epidermal autofluorescent molecules^10^ are the origin of the AF increases. Since cleavage of collagen by various proteases can lead to increased AF^43^, we proposed that the inflammation and oxidative stress inducers may increase the green AF by the following mechanisms: The inducers induce cleavage of full-length K1, leading to exposures of the K1’s domains that generate the AF. Our observation that NAC blocks both the UVC-induced K1 decrease and the UVC-induced AF increase is consistent with this proposal. It is of critical significance to further investigate the mechanisms underlying the AF increases.

Our study has not only provided first information indicating profound biomedical value of keratin’s AF, but also discovered a novel property of keratins’ AF: In addition to its basal AF, K1’s AF is inducible by inflammation and oxidative stress inducers. Moreover, our study has indicated that the green AF is a biomarker of ‘real damage’ of cells and tissues, since the intensity of LPS-, BSO-, or UVC-induced green AF is significantly associated with the dosages of these pathological insults that can dose-dependently produce cellular and tissue damage. In contrast, the current approaches for evaluating the risk of developing AIS can provide only mathematical probability of developing the disease based on the risk factors a person has^44^.

Collectively, our findings have formed basis for establishing a novel, keratin’s AF-based biomedical imaging technology. This technology has multiple distinct merits for non-invasive screening and diagnosis of inflammation- and oxidative stress-associated diseases. This technology may also be used for non-invasive, economic and rapid evaluations of the levels of inflammation and oxidative stress in the body of natural populations, which are critically needed in health management. Moreover, our study has found multiple novel biological properties, e.g., the AF asymmetry of AIS patients. Future investigation into the mechanism underlying these observations may expose new and crucial biological and pathological mechanisms.

## Acknowledgment

The authors would like to acknowledge the financial support by two research grants from a Major Special Program Grant of Shanghai Municipality (Grant # 2017SHZDZX01) (to L.J. and W.Y.) and a Major Research Grant from the Scientific Committee of Shanghai Municipality #16JC1400500 (to L.J.) and #16JC1400502 (to W.Y.).

## Legends of Supplementary Figures

**Supplementary Fig. 1**. (A) Nine d after injection of the LLC cells into the lung of C57BL/6 mice, all of the mice developed lung cancer. N = 4 - 7. (B) Administration of the anti-inflammation drug indomethacin did not significantly affect the weight of the lung with lung cancer. N = 4 - 7. ##, *P* < 0.01 (Student t-test); *, *P* < 0.05; **, *P* < 0.01. (C) H&E staining of the skin showed that the thickness of the epidermis was approximately 25 - 30 μm, while the thickness of the stratum corneum was less than3 μm. N = 6. Scale bar = 20 μm.

**Supplementary Fig. 2**. (A) UVC irradiation produced increased green AF of B16-F10 cells 0.3, 1 and 6 h after the UVC irradiation. Scale bar = 20 μm. (B) UVC irradiation produced significant and dose-dependent increases in the green AF intensity of B16-F10 cells 0.3, 1 and 6 h after the UVC irradiation. N = 12. *, *P* < 0.05; **, *P* < 0.01; ***, *P* < 0.001. The data were collected from three independent experiments. (C) The AF intensity of B16-F10 cells was significantly associated with the UVC dosages 0.3, 1 and 6 h after the UVC irradiation. The data were obtained from one representative experiment out of three independent experiments. N = 4. The *P* values of the other two independent experiments were < 0.001. The r values of the other two independent experiments conducted 0.3 h after the UVC exposures were 0.832 and 0.977, respectively; and r values of the other two independent experiments conducted 1 h after the UVC exposures were 0.910 and 0.940, respectively. (D) The spectra of both basal and UVC-induced AF of C57BL/6 mice, ICR mice and nude mice were similar, reaching maximal AF intensity at 470 - 500 nm when the excitation wavelength was 800 nm under a two-photon fluorescence microscope. N = 3. The data are the representative of the spectrum of the skin’s AF of one mice’s ear for each strain.

**Supplementary Fig. 3**. The spectrum of the AF of the AIS patient’s skin was virtually identical as that of the healthy controls. N = 3.

**Supplementary Fig. 4**. (A) Immunostaining assays of the ear’s skin of C57BL/6 mice using anti- K1 antibody also showed that the polyhedral structure of the K1-positive regions matched the polyhedral structure of the LPS-, BSO- and UVC-induced AF. N = 3-5. Scale bar = 5 μm. (B) Western blot assays on K1 using normal denaturing conditions showed that in the samples of the ear’s skin, there was a distinct band at approximately 127 kDa, while the level of K1 monomers at the 67 kDa band was exceedingly low. When the samples were prepared in 8 M urea, Western blot assays on K1 showed that the 127 kDa band disappeared with appearance of both a 67 kDa and a 59 kDa band. N = 3. (C) When the samples were prepared in 8 M urea, Western blot assays on K10 showed an increased band at 60 kDa. N = 3. (D,E) Western blot assays showed that the laser-based K1 siRNA administration led to a significant K1 decrease. The assays were conducted at 18 - 24 h after the K1 siRNA administration. N = 7. *, *P* < 0.05. (F) Immunostaining assays showed that the laser-based K1 siRNA administration led to a decreased K1 level in the ears of C57BL/6 mice. The assays were conducted at 18 - 24 h after the K1 siRNA administration. N = 3 - 5. Scale bar = 20 μm. (G,H) H2O2 induced a significant K1 decrease in B16-F10 cells. N = 8. *, *P* < 0.05. The data were collected from two independent experiments. (I, J) UVC induced a significant K1 decrease in B16-F10 cells. N = 9. #, *P* < 0.05 (Student t-test). The data were collected from three independent experiments.

## Legends of Supplementary Table

**Supplementary Table 1. Neither the LPS dosage nor the epidermal AF intensity was significantly associated with the Log-transformed serum levels of multiple cytokines 3 d after LPS administration**. Three d after the mice were administered with 0.5 or 1 mg/kg LPS, the green AF intensity of the ear’s skin of the mice was determined. The Log-transformed serum levels of multiple cytokines were also determined. N = 4 - 6.

**Supplementary Table 2. Neither the LPS dosage nor the epidermal AF intensity was significantly associated with the serum levels of multiple cytokines 3 d after LPS administration**. Three d after the mice were administered with 0.5 or 1 mg/kg LPS, the epidermal green AF intensity of the ear’s skin of the mice was determined. The serum levels of multiple cytokines were also determined. N = 4 - 6.

