## Supplementary figures and images for "Keratin-Based Epidermal Green Autofluorescence is a Common Biomarker of Organ Injury"

### Supplemental Fig. 1

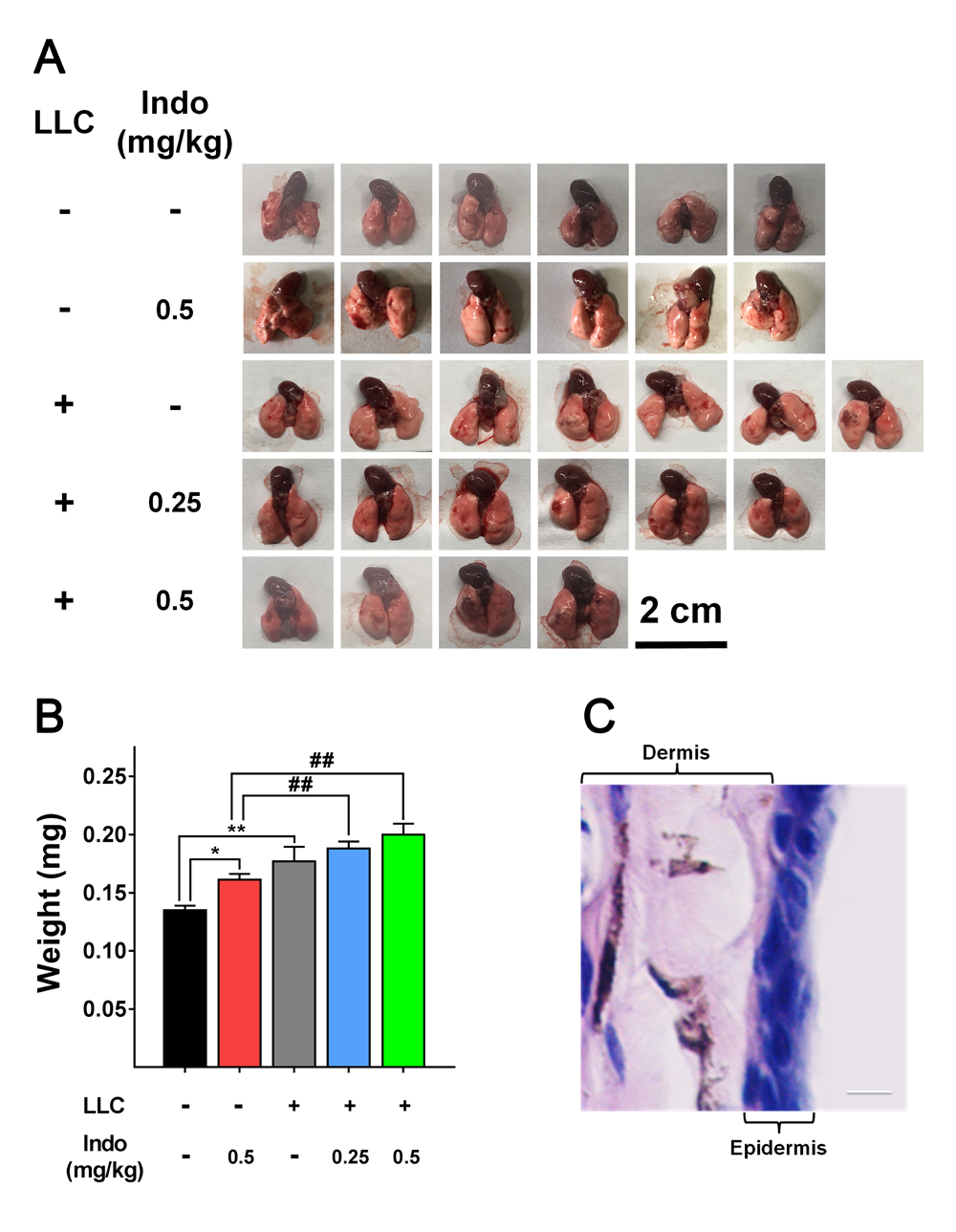

### Supplemental Fig. 2

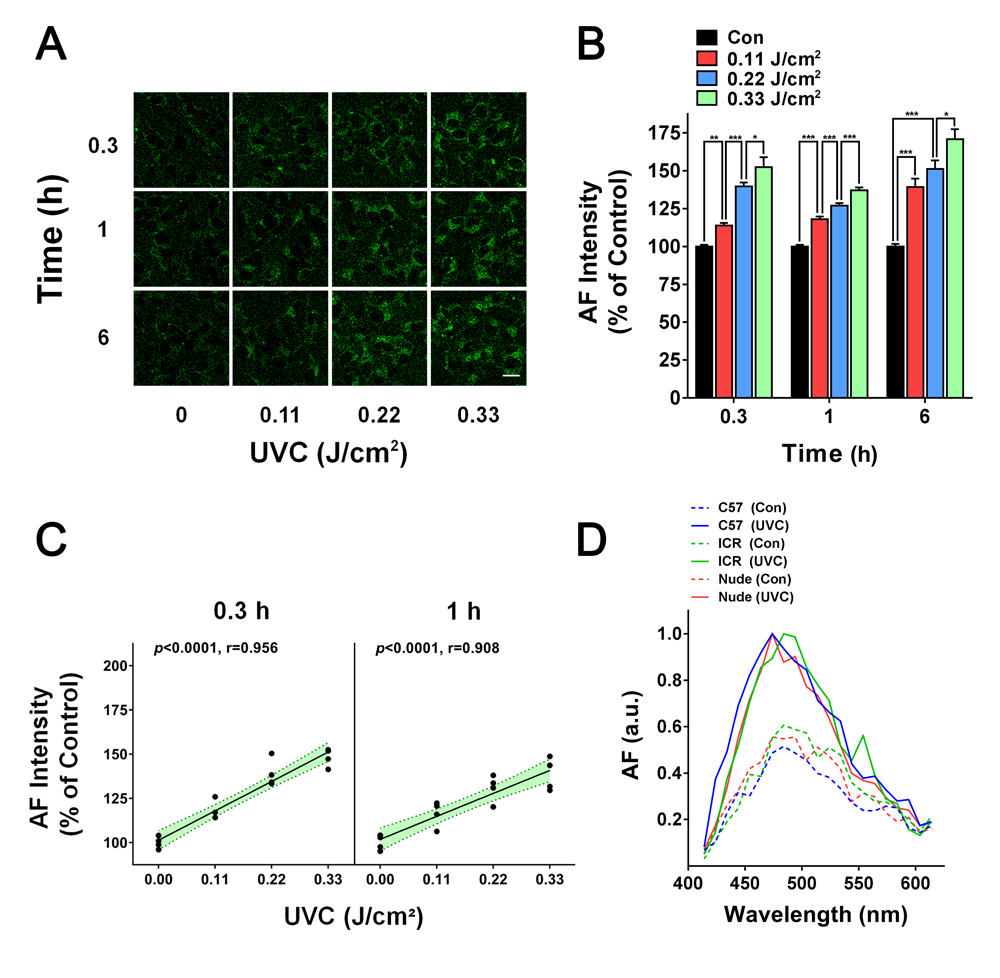

### Supplemental Fig. 3

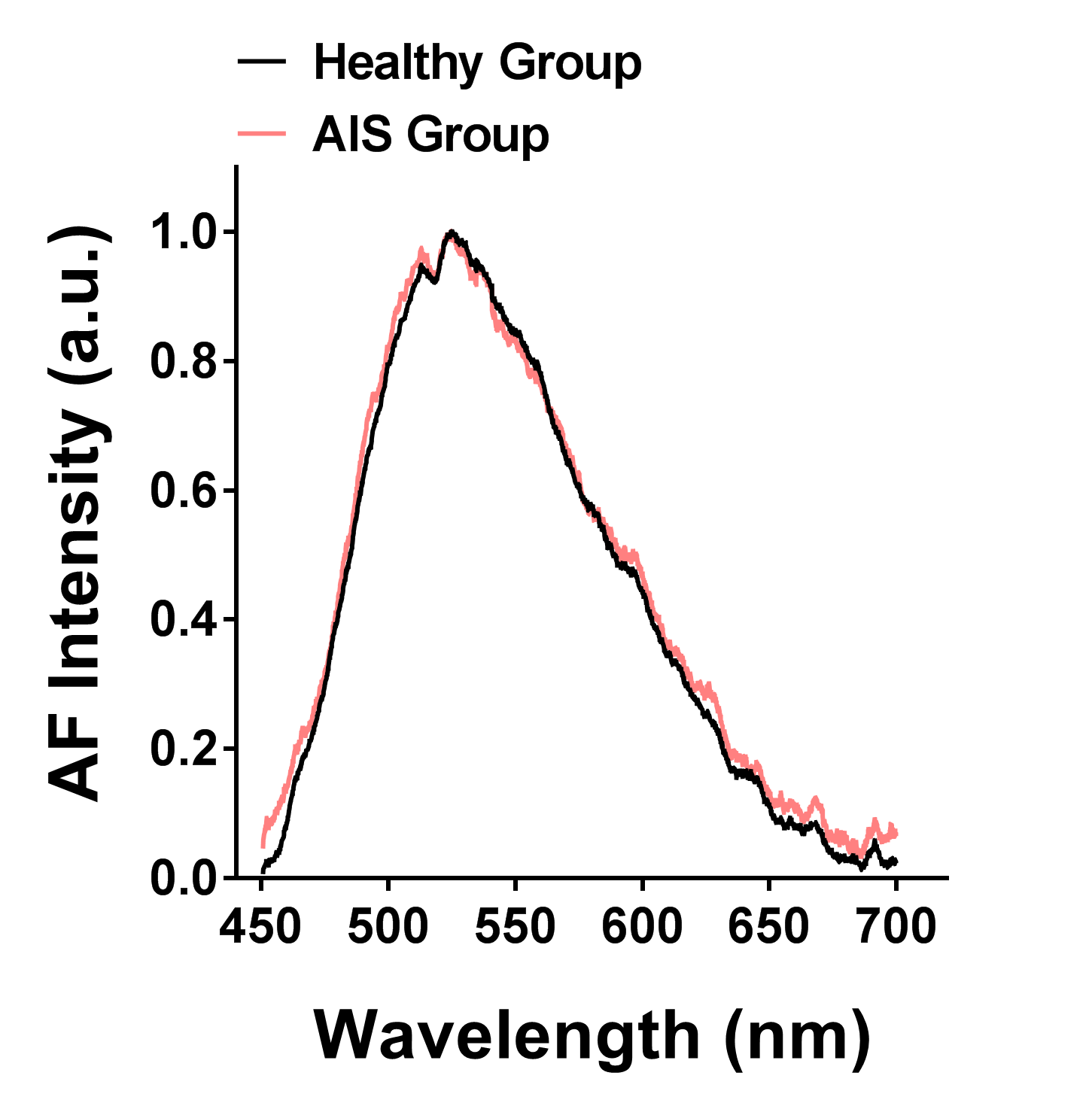

### Supplemental Fig. 4

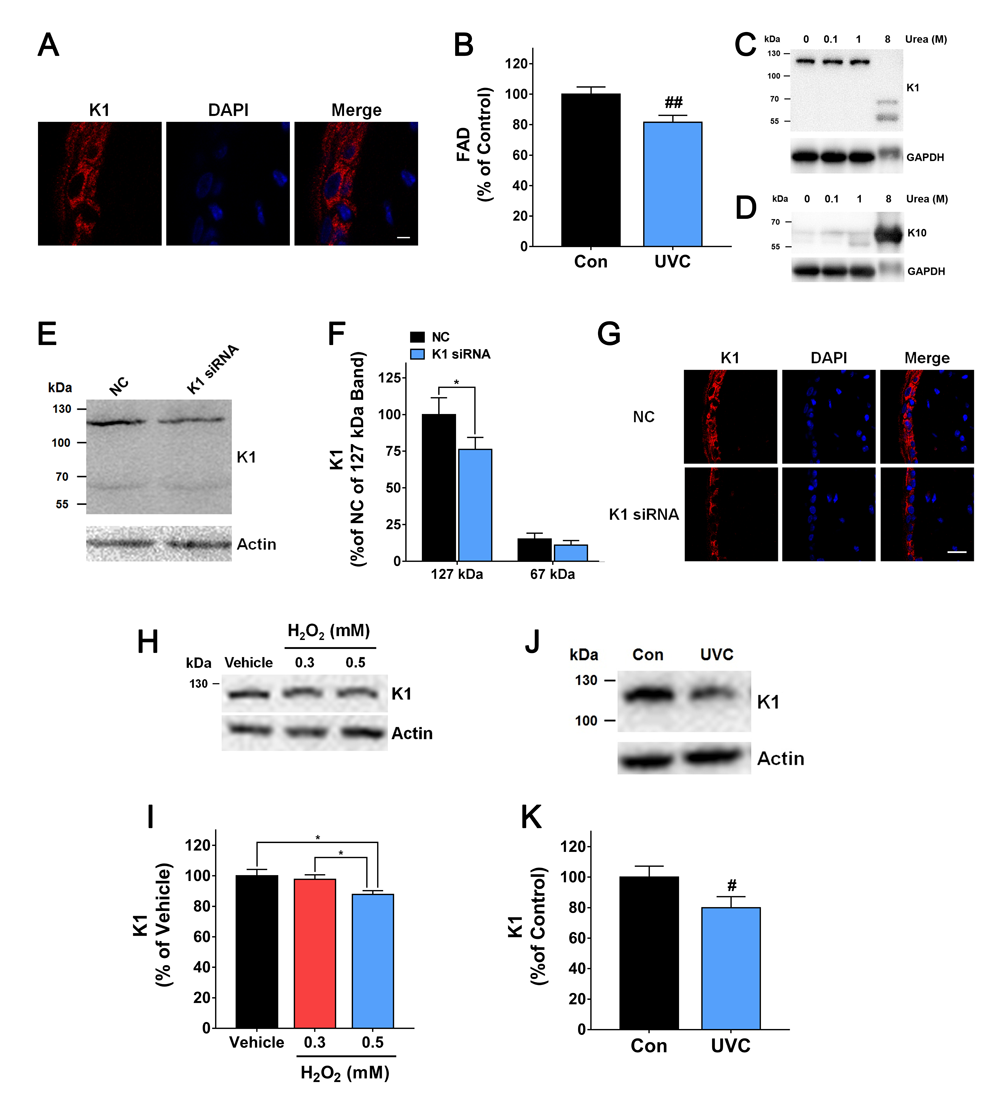

### Supplemental Table 1

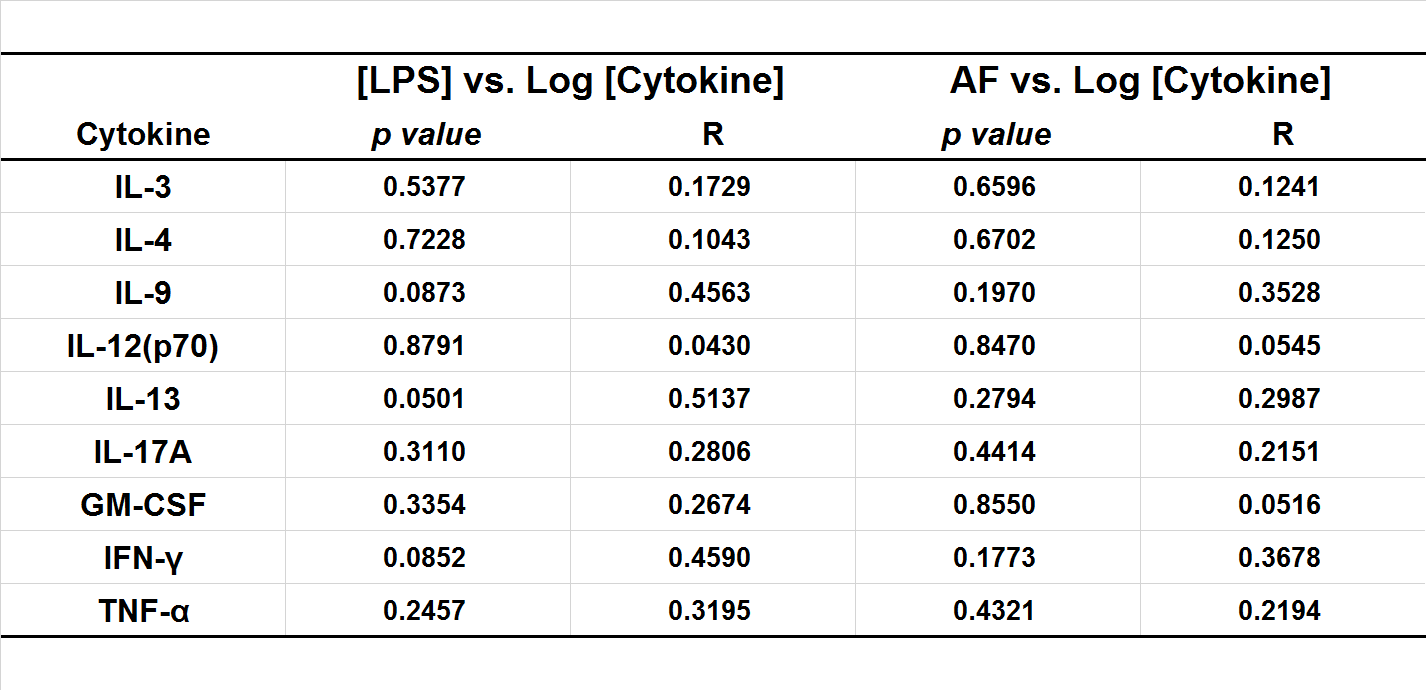

### Supplemental Table 2

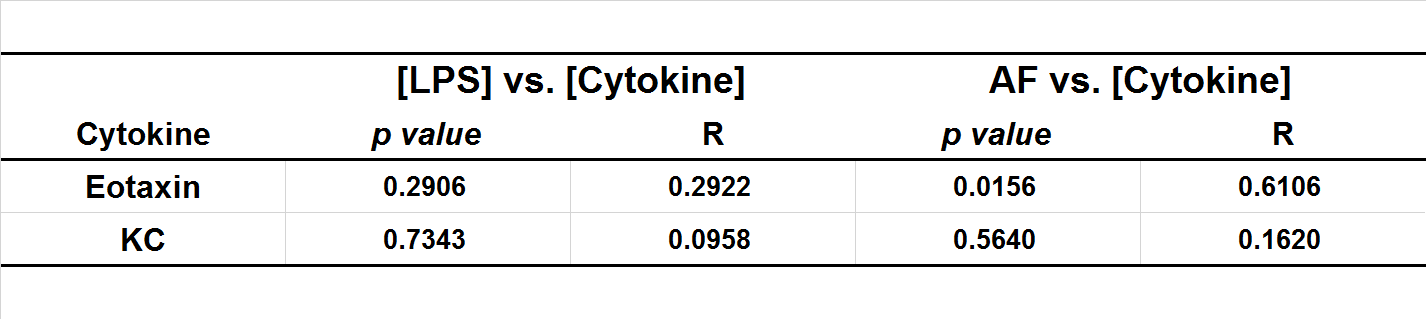
